# Cross-species variant-to-function analyses implicate *MEIS1* in conferring sleep abnormalities and impaired cerebellar development

**DOI:** 10.1101/2025.08.05.665539

**Authors:** Amber J. Zimmerman, Erika Almeraya Del Valle, Matthew C. Pahl, Fusun Doldur-Balli, Brendan T. Keenan, Patrick Z. Liu, Zoe Y. Shetty, Trisha R. Tsundupalli, Justin Palermo, Anitra Krishnan, James A. Pippin, Andrew D. Wells, Olivia J. Veatch, Alessandra Chesi, Philip R. Gehrman, Alex C. Keene, Allan I. Pack, Struan F.A. Grant

## Abstract

Genome wide association study (GWAS) reports substantially outpace subsequent functional characterization. Pinpointing the causal effector gene(s) at GWAS loci remains challenging given the non-coding genomic residency of >98% of these signals. We previously implicated effector genes at GWAS loci for the complex and polygenic disorder of human insomnia using a high-resolution cell-specific, chromatin capture-based variant-to-gene mapping protocol, paired with sleep phenotyping in *Drosophila*. In this study, we leveraged a diurnal vertebrate model with higher genomic conservation, namely zebrafish, to screen our six highest confidence candidate genes and identify those whose loss-of-function impaired sleep characteristics related to human insomnia-like behaviors. Of these genes, we observed that CRISPR-mediated deletion of the zebrafish ortholog of *MEIS1* produced nighttime specific sleep fragmentation and increased latency to sleep, pointing to a conserved role for *MEIS1* in sleep maintenance. Comparing our human cell-based chromatin accessibility and contact maps with publicly available zebrafish spatial genomic data revealed highly conserved genomic architecture harboring the insomnia GWAS variant of interest. Notably, this genomic conservation was selective for the zebrafish ortholog which contributed to the sleep phenotype, *meis1b,* while the duplicated ohnolog *meis1a* proved dispensable. Motivated by this, we characterized the spatio-temporal expression of *meis1b* in zebrafish, showing it is comparable to human with respect to cerebellar granule progenitors. Ultimately, we found that loss of *meis1b* impairs cerebellar development. Together, our work provides a powerful model for screening human disorder risk genes for sleep fragmentation using a tractable vertebrate and supports a conserved cerebellar role for *MEIS1* in sleep disturbance.

## Introduction

Nearly one-third of the adult population reports chronic sleep disturbance and symptoms of insomnia^1^, yet the underlying biological mechanisms remain largely uncharacterized. Insomnia represents a combination of difficulty initiating sleep (increased sleep latency) and/or difficulty maintaining sleep often accompanied by daytime consequences (e.g. fatigue, irritability) despite ample opportunity for sleep^2,3^. Insomnia, along with other sleep traits such as sleep duration, napping, and daytime dozing, is heritable and highly polygenic^4,5^.

Large-scale genome-wide association studies (GWAS) have revealed several hundred loci for insomnia and other sleep traits, the majority of which reside in the non-coding portion of the human genome, obscuring how they mechanistically confer their effect on susceptibility to a given trait^4–8^. Despite efforts to identify the actual causal “effector” gene(s) at each of these GWAS loci^4–8^, the majority of such non-coding signals remain simply either annotated to the nearest gene or mapped based on *in silico* prediction^6,7^. These approaches can misidentify the true effector gene(s) at a given GWAS locus^9–14^, which in turn can mislead the understanding of mechanisms underlying common disorders such as insomnia. Ultimately, this limits the utility of human genomic data for investigation of disease mechanisms and in driving forward therapeutic development.

Human cell-based fine-mapping of genome-wide loci associated with trait variation represents a targeted method to implicate disease-relevant effector genes^15–20^. Such leads can then be assessed through subsequent functional phenotyping in model organisms^16,21,22^ to characterize targets and their mechanistic role in disease etiology.

Our previous work nominated six high-confidence insomnia-associated genes (*SKIV2L*, *GNB3*, *CBX1*, *MEIS1*, *TCF12*, and *ARFGAP2*) through the integration of insomnia GWAS findings with variant-to-gene mapping in our human neural progenitor cell chromatin capture-based datasets^16^. Subsequent neuron-specific RNAi-mediated knockdown of these genes impacted sleep duration in *Drosophila*^16^. To extend our mechanistic interpretation of these novel genes, we sought to next employ a vertebrate model system, zebrafish, with higher genetic and cellular conservation relative to humans. This established and genetically tractable vertebrate model organism allows for a rapid developmental timeline that can aid efficient CRISPR-Cas9 mutagenesis paired with large-scale sleep and molecular phenotyping^22,23^.

Unlike most rodent models, sleep in zebrafish is diurnal and primarily consolidated to the night^24^. Additionally, zebrafish sleep is circadian-regulated, reversable, and has a heightened arousal threshold, making it an appropriate model for assaying multiple conserved traits relevant to sleep dysfunction^22,25–27^. Furthermore, much of the genomic organization between zebrafish and humans remains intact for many of the critical regulatory regions marked by syntenic conservation across vertebrates^28^. This syntenic conservation offers an opportunity to identify highly conserved genomic regulatory elements, such as enhancers or distal-interacting regulatory regions^29^, which frequently harbor GWAS-implicated non-coding variants. Leveraging these features, we sought to assess zebrafish sleep patterns following loss of these six nominated genes from our prior study and conduct subsequent mechanistic follow-up work focused on the highly conserved regulatory region around the most promising candidate from our screen, *MEIS1*.

The *MEIS1* locus is among the most statistically significant insomnia GWAS signals^4,6–8^. Though, there has been substantial debate as to whether its role in sleep disturbance is primarily driven by its association with restless leg syndrome (RLS) and/or with periodic limb movements in sleep (PLMS)^6,30–36^. These traits share an association signal residing within intron 8 of *MEIS1*, which has been shown to harbor an enhancer involved in the development of the caudate/basal ganglia^32,37^; thus, the majority of the work on *MEIS1* in this context has been focused toward this brain region. However, and importantly, our prior nomination of *MEIS1* from our variant-to-gene mapping was observed through a physical promoter-interacting chromatin contact^16^ with an additional independent signal located 5’ to *MEIS1*^7^, tagged by proxy rs13033745. This association is reported as being independent of the RLS or PLMS associations^31^. Therefore, we went on to characterize the biological relevance of this relatively overlooked signal at this key locus. Together, we performed a thorough investigation of the *MEIS1* region to assess the underlying causal mechanism(s) at this key insomnia locus.

## Materials and Methods

### Animal use

All experiments with zebrafish were conducted in accordance with the Institutional Animal Care and Use Committee guidelines under the appropriate institutional approved protocols (806646 and IAC 21-001154) at the University of Pennsylvania and the Children’s Hospital of Philadelphia. Breeding pairs consisted of wild-type AB and TL (Tupfel Long-fin) strains. Fish were housed in standard conditions with 14-hour:10-hour light:dark cycle at 28.5°C, with lights on at 9am (ZT0). and lights off at 11pm (ZT14).

### CRISPR/Cas9 mutagenesis

Highly specific guide RNAs (gRNAs) were designed using the online tool Crispor (http://crispor.tefor.net/) with the reference genome set to “NCBI GRCz11” and the protospacer adjacent motif (PAM) set to “20bp-NGG-Sp Cas9, SpCas9-HF, eSpCas9 1.1.” gRNAs were prioritized by specificity score (>95%) with 0 predicted off-targets for sequences with up to 3 mismatches. The zebrafish sequence was obtained using Ensembl (https://useast.ensembl.org/) with GRCz11 as the reference genome. Sequence was aligned to the human amino acid sequence using MARRVEL (http://marrvel.org/) and DIOPT to identify the region with highest conservation, and each gRNA was designed targeting the most 5’ conserved regulatory region that was present in all transcripts. Exon 1 was skipped to avoid usage of potential alternative start codons^38^. For *meis1a*, two gRNAs were designed because one was not sufficient to disrupt all transcripts. gRNA sequences are provided in **Table S1**. AB/TL breeding pairs were set up overnight and embryos collected in embryonic growth media (E3 medium; 5mM NaCl, 0.17 mM KCl, 0.33 mM CaCl2, 0.33 mM MgSO4) the following morning shortly after lights-on. Pre-formed ribonuclear protein (RNP) complexes containing the gRNA:tracr and spCas9 HiFi enzyme were injected at the single-cell stage alternating between the gene group and scramble-injected sibling control group. Injections of scramble gRNA and gene-specific gRNA were performed in embryos from the same breeding event (clutch). Embryos were left unperturbed for one day before being transported to fresh E3 media in petri dishes (approximately 50 per dish).

### DNA extraction and PCR for genotyping

DNA extraction was performed per the manufacturer’s protocol (Quanta bio, Beverly, MA) immediately following completion of the sleep assay, as described previously^16^. Larvae were euthanized by rapid cooling on a mixture of ice and water between 2-4°C for a minimum of 30 minutes after complete cessation of movement was observed. Genotyping was performed on individual fish at the conclusion of each sleep assay. Either restriction digest or headloop PCR methods were used to validate mutations and confirm editing efficiency^16,39,40^. Headloop primers were designed to overlap the Cas9 cut site so that mutation of this site would block headloop formation leading to amplification of the target sequence^39,40^. Otherwise, gRNAs were designed to disrupt a restriction enzyme site such that effective mutation could be detected by a lack of restriction enzyme cutting. For headloop primers, mutation efficiency was calculated as the ratio of the headloop amplicon: standard primer amplicon band intensity using ImageJ^16,40^. For standard primers, mutation efficiency was calculated as the ratio of the uncut:cut amplicon. Phenotype inclusion criteria was >90% mutation efficiency. Select samples were Sanger sequenced to confirm the on-target Cas9 cut 3-4 bases upstream of the Protospacer Adjacent Motif (PAM). Mutant sequences were aligned to negative controls using SnapGene to show the consequence of the mutation. Out-of-frame mutations were considered those that produced base changes in non-multiples of 3^38^. Primers for genotyping and restriction enzymes are listed in **Table S2**. All primers were run on a 2% agarose gel and sequence verified using Sanger sequencing to verify the target region was absent of polymorphisms that may hinder genomic editing.

### Data collection and analysis for sleep phenotyping

All embryos and larvae were housed in an incubator at 28.5°C, with lights on at 9am (ZT0) and lights off at 11pm (ZT14) prior to data collection. Dead embryos and chorion debris were removed daily until day 5 post fertilization. On day 5, CRISPR mutants and scramble-injected sibling controls were screened for gross morphological deficits and healthy larvae were pipetted into individual wells of a 96-well plate and placed into a Zebrabox (ViewPoint Life Sciences) for automated video monitoring. Genotypes were placed into alternating rows to minimize location bias within the plate. All animals were allowed to acclimate to the Zebrabox for approximately 24 hours before beginning continuous data collection for 48 hours starting at lights on (9am). At least two biological replicates were run for each gene using different clutches (sibling-matched within clutch) of embryos and well placement was flipped for each experiment to minimize location bias across experiments. Each Zebrabox is sound-attenuating and contains circulating water held at a temperature of 28.5°C with automated lights cycling on the same 14-hour:10-hour light/dark cycle. Activity data were captured using automated video tracking (Viewpoint Life Sciences) software in quantization mode. As described previously^16,41^, threshold for detection was set as the following: detection threshold: 20; burst: 29; freeze: 3; bin size: 60 seconds. Data were processed using custom MATLAB scripts^42^ to calculate the following parameters for both day and night separately: sleep duration (minutes/hour), activity duration (seconds/hour), waking activity (seconds/awake minute/hour), sleep bout length (minutes/bout), sleep bout number (number/hour) and nighttime sleep latency (minutes).

### Arousal Threshold Assay

A mechano-acoustic stimulus was used to determine arousal threshold using a protocol adapted from previous work^43,44^. Individual fish were placed into alternating columns of a 96-well plate to avoid location bias. Ten different vibration frequencies were applied, which consistently produced a step-wise increase in arousability. Frequencies were pseudo-randomly assigned to prevent acclimation to any given stimulus frequency throughout the trials. Frequency steps of 40Hz were ordered as follows: 560Hz, 400Hz, 520Hz, 720Hz, 440Hz, 680Hz, 480Hz, 760Hz, 600Hz, and 640Hz. These ten frequencies were each presented ten times for 5 seconds every 3 minutes (i.e. 5 seconds on, 2 mins 55 seconds off) beginning at 1am (ZT16) and ending at 6am (ZT21). Lower frequencies produced larger changes in movement and an increase in the fraction of responsive larvae. Therefore, analyses were presented as highest-to-lowest frequency representing lowest-to-highest relative intensity of stimulation (i.e., 760Hz, 720Hz, 680Hz, 640Hz, 600Hz, 560Hz, 520Hz, 480Hz, 440Hz, and 400Hz). The response to stimuli was measured by automated video tracking and analyzed using MATLAB (Mathworks)^42^ and Excel (Microsoft). The response fraction represents the percentage of larvae considered to be asleep during the 5 second pre-stimulus baseline (activity <0.01s/5sec, determined empirically using average movement prestimulus) and whose activity increased over baseline during the stimulus presentation. Pre-stimulus activity is calculated as average activity for each fish (n = 29 negative control, 32 *meis1b* crispants) prior to all stimuli (100 stimuli total). Proportion of fish awake prior to each stimulus is calculated as the total number of fish awake (activity <0.01s/5sec) prior to each stimulus. The proportion is calculated for each stimulus (100 separate pre-stimulus events), thus the sample size for proportion awake is 100 in each group. Curve-fit and statistical analyses were carried out in Prism V9 (Graphpad) using non-linear regression (Agonist vs Response--Variable slope [four parameters]) with extra sum-of-squares F-test used to compare EC50 (half-maximal response) and top (maximal response) as described previously^43^. EC50 effectively represents the arousal threshold for the population of fish.

### Statistical analysis and control for multiple comparisons across sleep traits

Sample size determination was based on previous work^40^, indicating 48 larvae per group (half of one 96-well plate) is sufficient to detect medium effects at a power of 0.8 and significance level of 0.05 even in a mixed population of mutants. Experiments were repeated in the same Zebrabox at least twice. Effect sizes are described as standardized mean differences (SMDs) and 95% confidence intervals (CI). Phenotypes of interest included a total of eleven measurements related to sleep and activity, including day and night sleep duration, activity, waking activity, sleep bout length, and sleep bout number, as well as nighttime sleep latency. Analyses compared these traits between scramble-injected and CRIPSR mutant (crispant) fish. Fish with more than one outlier value on an activity metric (±2 SD) were removed from analyses on all parameters. Primary comparisons between crispant and negative control-injected siblings were performed using non-parametric Wilcoxon rank-sum tests to mitigate any potential impact of non-normality of the endpoints. A Hochberg step-up procedure^45,46^ was applied independently to analysis of each gene to maintain gene-specific type I error at the desired level of 0.05 across the multiple phenotypes being assessed. Statistical analyses and experimental numbers for activity monitoring are provided along with a README file through DRYAD. http://datadryad.org/share/I07vjxcKg3i5Atl_H1A60C02e3A-gFLA2bLXofNpC7E.

### Variant-to-gene mapping

Details regarding ATAC-seq, RNA-seq, and Capture C protocols along with variant and loop calling are described in the initial publication detailing candidate effector gene identification^16^. In addition to human induced pluripotent stem cell (iPSC)-derived neural progenitor cells (NPC), we performed the same loop calling procedure specifically for rs13033745 in human primary astrocytes and human iPSC-derived neurons.

### Whole mount *in situ* hybridization chain reaction

Larvae used for HCR were treated with PTU (0.003%) between 22-24 hours post fertilization to block pigment and were maintained in embryo media containing PTU (0.003%) until euthanasia. Larvae were fixed in 4% PFA at 4 days post fertilization for approximately 24 hours then washed with 1X PBS 3 times for 5 minutes. Whole larvae were washed for 5 minutes 3 times with 1X PBST. *In situ* hybridization protocol was performed as previously described^47–49^. Custom probes were designed using amplifier sequences previously described^49^ to generate 30 probes (15 pairs) for *meis1b*, *neurod6b,* and *pax6a*. Forty probes (20 pairs) were designed for *meis1a* due to predicted lower expression in brain. Probe design was performed using publicly available code from github^50^ https://github.com/rwnull/insitu_probe_generator. HCR buffers and B amplifiers were purchased from Molecular Instruments (https://www.molecularinstruments.com/). B2-488 was used for *pax6a*, B3-546 was used for *neurod6b*, and B4-647 was used for *meis1b* and *meis1a*.

### Bodipy staining

Bodipy^493/503^ was used to stain the cerebellar neuropil. Zebrafish larvae were raised to 4 or 6 days post fertilization then euthanized and fixed in fresh 4% PFA for 24 hours. Whole larvae were then washed in 1X PBS 3 times for 5 minutes followed by one time in 1X PBST for 5 minutes. Four-to-six larvae were placed into 500μl of 1xPBS with 0.5μl (1:1000) of Bodipy for approximately 15 minutes prior to mounting.

### Confocal imaging

Whole zebrafish larvae were mounted in 1% low melting agarose against Corning glass coverslips No. 1.5 and imaged using a Zeiss LSM 980 Confocal microscope. For HCR imaging, laser settings were held consistent for every image using the following parameters: 647 at 60% power, 600V gain; 568 at 25% power, 650V gain; 488 at 25% power, 600V gain. For Bodipy imaging, laser settings were held at the following: 488 - 4% power, 600V gain. Images of whole larvae were captured using a 5x objective (videos show z-stacks at 10x). Images captured for cerebellar analysis were captured using 10x objective. Z-stacks were taken at 1µm intervals for whole larvae and 5µm intervals for cerebellar images with the following parameters: 1024X1024 pixels, line averaging-2x for HCR, 4x for Bodipy, bidirectional scan, 8-bits per pixel. Confocal z-stacks were visualized using FIJI (ImageJ). Single z-slice images were created for each image used to examine whole body expression. Three-dimensional projections were created to measure cerebellum and upper rhombic lip to capture full width of cerebellum and upper rhombic lip.

### Public data curation and analysis

UCSC tracks for conservation and GTEx eQTLs, were obtained within the UCSC genome browser using hg19 coordinates. Histone marks including H3K27ac and H3K4me3 for neurons and PLAC-seq for oligodendrocytes were obtained from Nott et al., 2019^51^. TAD boundaries for h1-NPCs^52^ were downloaded from the 3D genome browser (http://3dgenome.fsm.northwestern.edu).

Linkage disequilibrium *r*^2^ values were computed using LDLink with the LDProxy tool against GRCh37 (hg19) with the ancestry set to CEU and a calculation window of 500Kb.

WashU genome browser datasets for zebrafish brain were obtained from Yang et. al., 2020^53^ provided in danRer10 coordinates. Data are expressed as fold change and brain cell type 8 was used for scATACseq from adult zebrafish brain as it is representative of atoh1-expressing (granule progenitor) cells.

GTEx eQTL data from the cerebellum were obtained for rs13033745 from https://www.gtexportal.org.

Single cell datasets were obtained from the following resources for human cerebellar development (Sepp et al., 2024^54^ available at https://apps.kaessmannlab.org/sc-cerebellum-transcriptome/) and zebrafish development^55^ (https://daniocell.nichd.nih.gov). Data were visualized using *Seurat* in R.

## Results

### Screening of candidate genes for insomnia-like behaviors in zebrafish implicates meis1b

We employed CRISPR-Cas9 mutagenesis at the single cell-stage of zebrafish^40^ development to target each of the six high-confidence candidate genes (*SKIV2L, GNB3, ARFGAP2, MEIS1, TCF12,* and *CBX1*)^16^ and assayed 11 sleep-wake behaviors using automated video monitoring in both F0 CRISPR mutant (crispant) larvae and scramble-injected sibling controls aged 5-8 days post fertilization (**Figure 1A**). We selected orthologues to the six genes resulting in nine genes, given the genomic duplication of *gnb3*, *cbx1*, and *meis1*. We designed guide RNAs that targeted critical exonic domains to produce predicted loss-of-function mutations and then confirmed that the on-target mutations produced out-of-frame mutations for each guide RNA (**Figure S1**).

**Figure 1.**
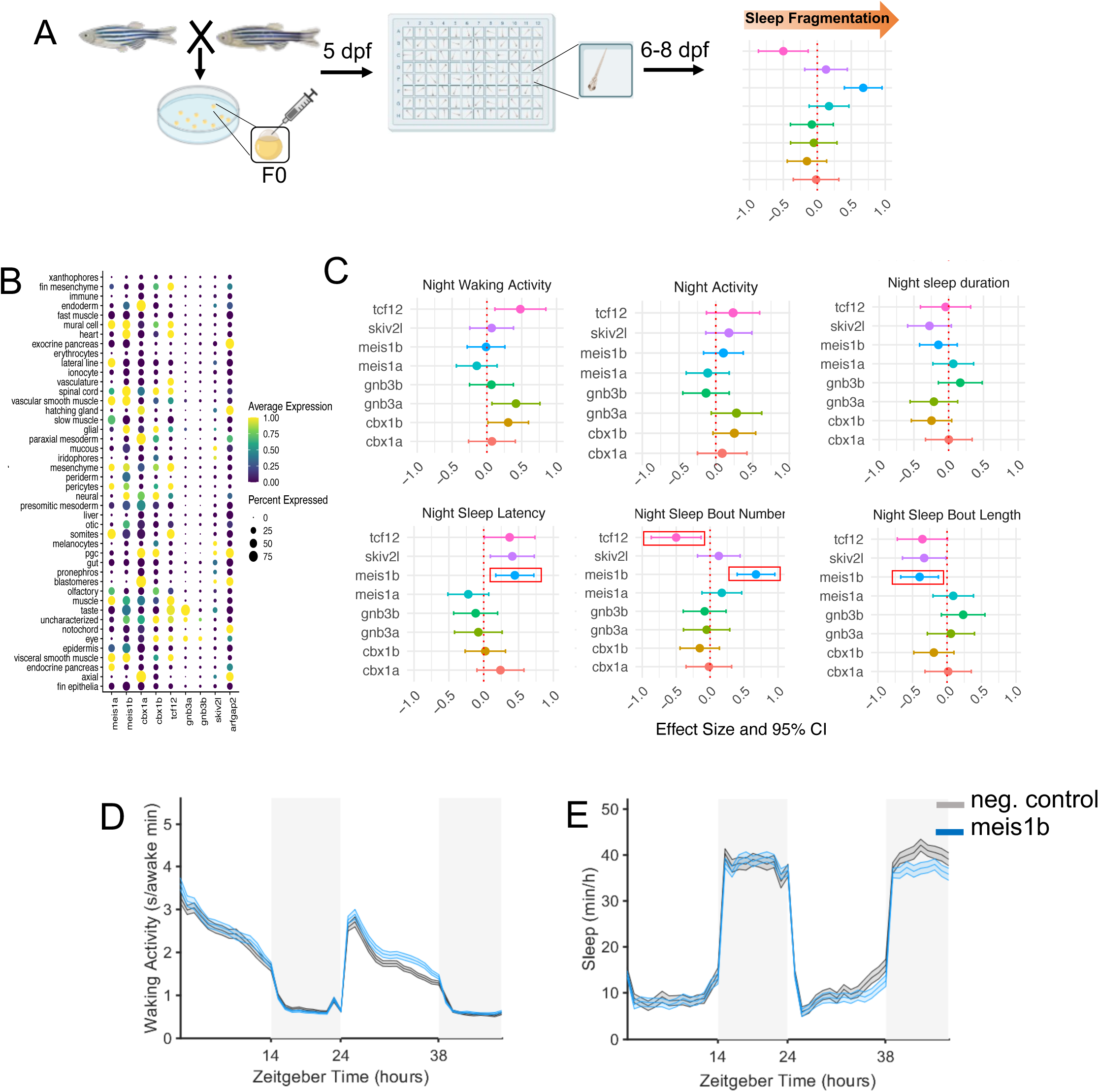
Zebrafish screen of insomnia-associated genes. A. Paradigm of screening approach. Wild-type AB/TLF embryos are injected at the single cell stage with preformed ribonuclear protein complexes containing Cas9 and tracr:cRNA to selectively mutate conserved domains of each gene. At 5dpf, larvae are transferred to individual wells of a 96-well plate and continuous activity is captured from 6-8dpf to measure sleep and activity parameters. B. Scaled expression of the orthologous candidate insomnia genes during larval zebrafish development (data from Daniocell). C. Selected sleep/activity parameters associated with human insomnia-like behaviors. D. Mean ± SEM waking activity in *meis1b* crispants on days 6-7dpf. E. Mean ± SEM sleep in *meis1b* crispants on days 6-7dpf. In C, sleep data are presented as effect size (standardized mean difference) ±95% confidence interval (CI) for each crispant group compared to its own sibling-matched controls. Data are pooled from 2 independent experiments. N for each experiment: *skiv2l* (80 crispants/ 77 controls), *meis1a* (94 crispants/ 86 controls), *meis1b* (111 crispants/ 100 controls), *gnb3a* (68 crispants/64 controls), *gnb3b* (75 crispant/ 80 controls), *cbx1a* (60 crispant/ 79 controls), *cbx1b* (102 crispant/ 83 controls), (74 crispant, 49 controls), *arfgap2* (58 crispant/79 controls). Red boxes indicate measures that reached significance following correction (see **Methods**).

To ensure each of these genes was expressed at the larval zebrafish stage when we tested sleep, we first reanalyzed public domain single cell RNA sequencing (scRNAseq) data from zebrafish across development for a broad range of cell types^55,56^, showing that the genes encoding transcription factors (orthologs to *MEIS1*, *CBX1*, and *TCF12*) are most abundantly expressed, but expression varies widely across cell types (**Figure 1B**). The duplicated ohnologs do not show concordant expression patterns, a common feature in zebrafish where duplication of genes results in cell- and temporal-specific expression for each^57^. Given this, we elected to target each ohnolog independently to identify phenotypes that could be driven by specific cell types where each is expressed.

To identify crispants in our experimental setting which produced relevant behavioral phenotypes, we defined *insomnia-like* behavior in zebrafish as an increase in latency to sleep onset at night paired with fragmentation of nighttime sleep measured as an increase in number of sleep bouts. Loss of *arfgap2* impaired development of the notochord and motor function (as reflected by dramatically reduced waking activity) (**Figure S2B-C)**. This phenotype is reflective of its high expression in the notochord (**Figure 1B**); thus, we removed this crispant from our subsequent grouped sleep analyses. Each ohnolog (e.g. *meis1a* vs. *meis1b*) was tested separately, and we observed discordant behavioral patterns for duplicated pairs across most metrics (**Figure 1C** and **Figure S2A**), reflecting the cell-specificity of each ohnolog observed in the scRNAseq data (**Figure 1B**).

Consistent with our previous work in *Drosophila*^16^, loss of *skiv2l* produced the largest effect on sleep duration (**Figure S2A**). Mutation of *meis1b* produced a behavioral response within the *a priori* defined phenotype related to “insomnia-like,” based on significantly increased latency to sleep onset, along with more frequent and shorter sleep bouts at night and no effect on general waking activity (**Figure 1C-E**, **Figure S2A, and Table 1**). Conversely, mutation of *meis1a* did not significantly impact sleep measures (**Figure 1C, Figure S2A and Table 2**).

Given the known role of *MEIS1* in hindbrain development^58,59^ as well as its association with RLS^30–33^, we next tested for changes to motor activity using a variety of activity metrics and arousal threshold, which measure depth of sleep in response to increasing stimuli intensities (**Figure S3A-B**). Despite having highly fragmented sleep, *meis1b* crispants did not present with heightened total activity at night (**Table 1 and Figure S3C-D**), nor did they have changes in arousal threshold (**Figure S3E**). Together, these data demonstrate *meis1b* plays a role in sleep-wake transitions rather than sleep depth or hyperactivity.

### The 5’ MEIS1 association implicates the cerebellum via cis-regulation of MEIS1-AS3

Of the independent insomnia signals established at *MEIS1*^6^ that are not in LD with the RLS association, only the 5’ variant, rs13033745, harbors a cis-regulatory element interacting with the *MEIS1* promoter in our data^16^ (see **Figure 2**); thus, we focused our functional interrogation on this variant.

**Figure 2.**
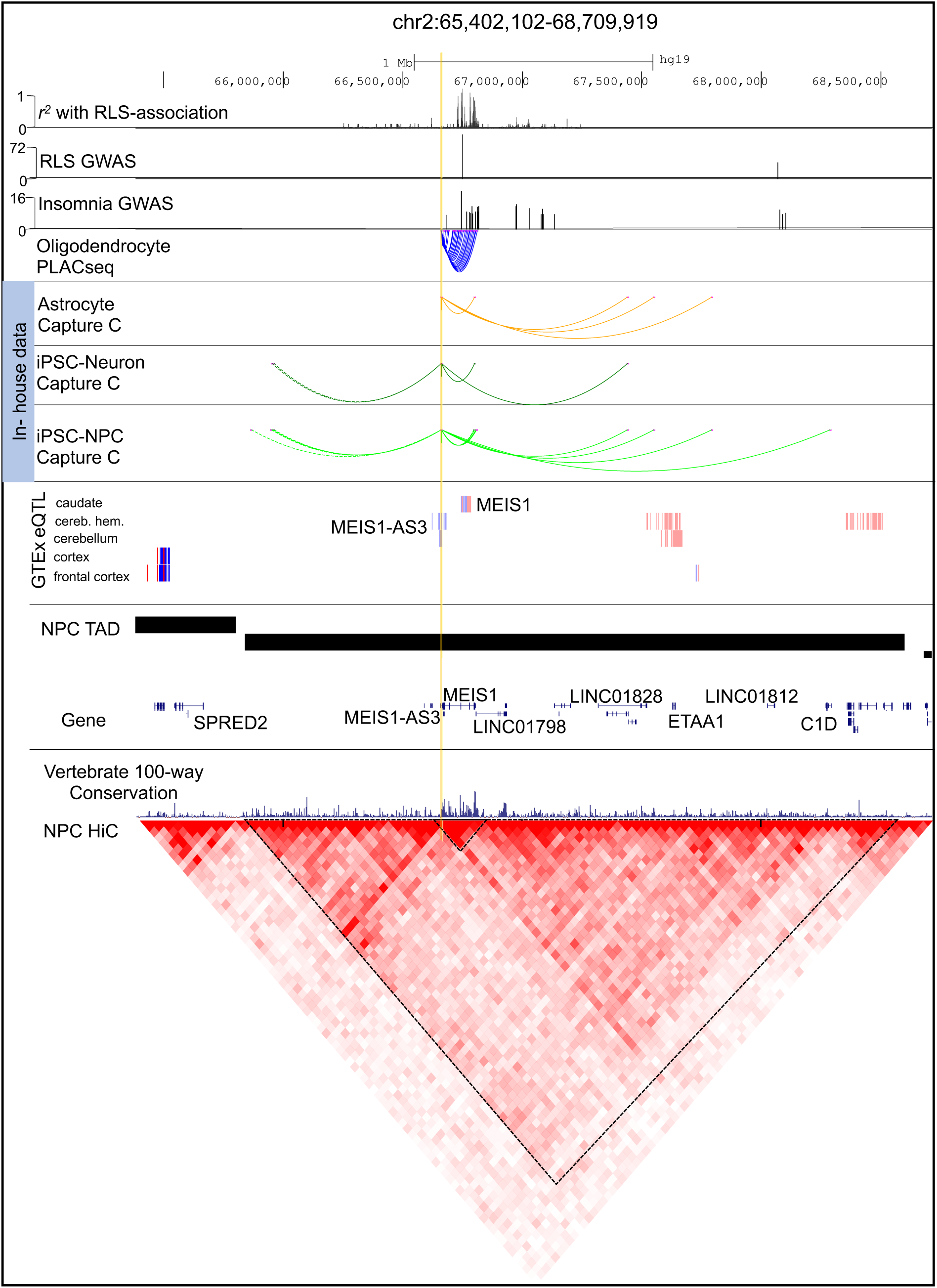
Variant-to-gene mapping at the *MEIS1* locus. UCSC genome browser plot showing the *MEIS1* region in hg19 coordinates. From the top: *r*^2^ values for the RLS haplotype tag SNP (rs113851554) that is in LD with the most significant association signal for insomnia in intron 8 of *MEIS1*. Independent RLS GWAS associations reported previously^36^. Independent insomnia GWAS associations reported previously^6^. The height of the bars representing GWAS signals indicates the z-score. Putatively causal variant we identified previously^16^, rs13033745 (yellow line) *r*^2^>0.7 with sentinel rs1519102. Oligodendrocyte PLACseq loops from Nott et al.^51^ that interact with rs13033745. Promoter-focused capture C data from our human cell-derived primary astrocytes, neurons, and neural progenitor cells (NPCs). Loops are only shown for contacts with rs13033745. NPC TAD coordinates were downloaded from the 3D genome browser from Dixon et al H1-NPCs^52^. Gene coordinates. GTEx eQTL track showing eQTLs for *MEIS1-AS3* in the cerebellum and *MEIS1* in the caudate. Conservation track displays 100-way vertebrate conservation. NPC Hi-C data were obtained from the 3D genome browser (https://3dgenome.fsm.northwestern.edu/).

To first identify all proximal and distal cis-regulatory contacts with rs13033745 in cell types that highly express *MEIS1*, we analyzed our promoter-focused Capture C along with previously published PLAC-seq data from iPSC-derived neural progenitors, mature neurons, astrocytes, and oligodendrocytes (oligodendrocyte data from Nott et al. 2019^51^) (**Figure 2)**. We examined a 2.76Mb window (chr2:65,840,001-68,600,000) corresponding to the annotated topological associated domain (TAD) boundary from NPCs^52^ encompassing *MEIS1*. rs13033745 contacts multiple non-coding RNAs within the TAD, revealing highly reproducible contacts with promoter baits at *MEIS1-AS3*, *MEIS1,* and the distal long non-coding RNA*, LINC01798* (**Figure 2**). Notably, these contacts span a region of high conservation that corresponds to a frequently interacting sub-TAD encompassing *MEIS1* (**Figure 2**). We did observe contacts between rs13033745 and the other independent intron 8 association region in oligodendrocyte PLAC-seq; however, no intronic contacts were observed in any other cell type for rs13033745. This specific observation was not due to differences in PLAC-seq and Promoter Capture-C methodologies because we also examined neuronal PLAC-seq^51^ and did not observe intronic contacts with rs13033745 (data not shown), indicating cell-specific chromatin contacts exist between rs13033745 and distal regions.

Given the previous eQTL analysis implicated the intron 8 signal impacts *MEIS1* expression in the caudate^60^, we similarly mined eQTL data from GTEx for our 5’ signal of interest at the *MEIS1* locus. Interestingly, rs13033745 colocalizes with an eQTL for the anti-sense RNA, *MEIS1-AS3* in the cerebellum, and all other variants in this region are associated only with *MEIS1-AS3* in the cerebellum and not other brain regions (**Figure 2**). Together, these results indicate the 5’ insomnia risk variant, rs13033745, is involved in cis-genomic regulation through multiple proximal and distal chromatin contacts spanning the *MEIS1* locus. Importantly, via eQTL analysis, this 5’ association implicates the cerebellum as opposed to the caudate, which is associated with the intron 8 signal.

### MEIS1 promoter-interacting cis-regulatory elements are only conserved for the zebrafish ortholog, meis1b

The topological interactions at this key locus demonstrate that the *MEIS1* promoter is under distal regulation by multiple non-coding cis-regulatory elements associated with insomnia variants. Previous work has shown that syntenic conservation across species frequently maintains TAD boundaries, keeping intact genomic regulatory blocks^61,62^. Therefore, we examined the genomic architecture in the syntenic block encompassing each zebrafish *meis1* ohnolog with the aim of identifying ‘highly conserved non-coding elements’ (HCNEs) that correspond to those at the human *MEIS1* locus.

We integrated our assay for transposable accessible chromatin sequencing (ATAC-seq) and promoter-focused capture C datasets from human iPSC-derived neurons and NPCs with publicly available genomic data from both human^51^ and zebrafish^53^ to compare the genomic architecture across species. Syntenic organization exists for both zebrafish ohnologs, *meis1a* and *meis1b*, with consistent flanking genes, *ETAA1* and *SPRED2*, (**Figure 3B and Figure S4**) corresponding to the TAD structure in human (**see Figure 2**).

**Figure 3.**
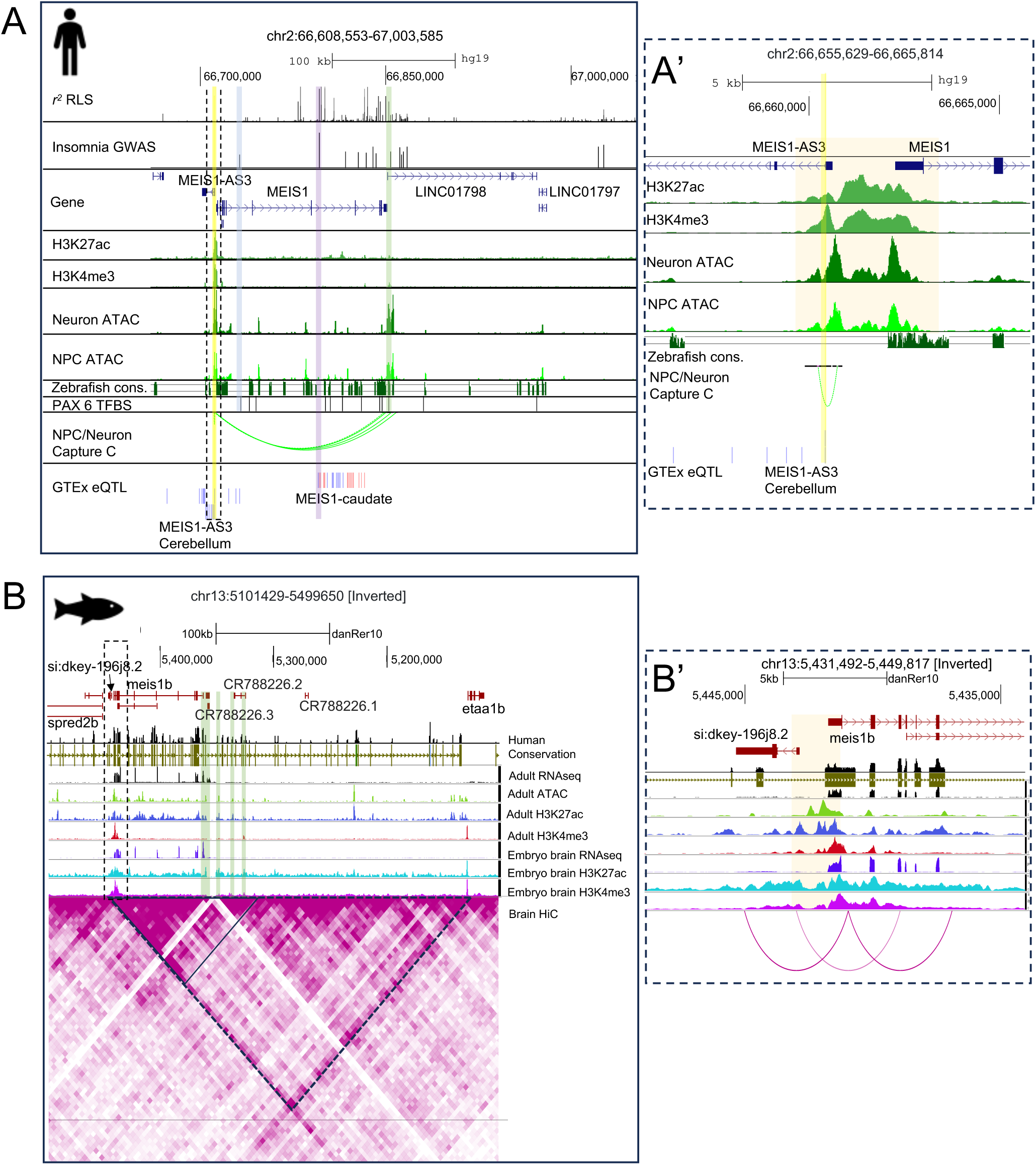
Comparison of human and zebrafish *MEIS1* genomic regions. A. UCSC genome browser plot of the highly conserved region around *MEIS1* displayed in Figure 2. From the top: *r*^2^ values for the RLS haplotype tag SNP (rs113851554) that is in LD with the most significant association signal for insomnia in intron 8 of *MEIS1* (purple highlight). Independent insomnia GWAS associations reported previously^6^. The height of the bars representing GWAS signals indicates the z-score. Putatively causal variant we identified previously^16^, rs13033745 (yellow highlight) *r*^2^>0.8 with sentinel rs1519102 (blue highlight). Gene coordinates. Histone marks H3K27ac and H3K4me3 obtained from Nott et al.^51^ ATACseq from our human iPSC-derived neurons and NPCs. Zebrafish conservation (green segments indicate regions of sequence homology) using Multiz alignment. Predicted PAX6 transcription factor binding sites (JASPAR). Note the overlap with distal promoter-interacting chromatin loop within *LINC01798* (shaded green bar). NPC/Neuron promoter-capture C loops for rs13033745. GTEX eQTLs. A’. Zoomed view of shared promoter region between *MEIS1* and *MEIS1-AS3.* rs13033745 is highlighted with a yellow bar. B. WashU epigenome browser plot showing chromatin landscape at the *meis1b* locus in zebrafish. Chromosomal coordinates are shown for the danRer10 genome build. Conservation is shown with reference to the human genome with human chain-net track colored by chromosome. Chromatin tracks show data from Yang et al.^53^ for RNAseq, ATACseq, H3K27ac, and H3K4me3 marks from zebrafish adult brain and embryo along with a HiC interaction plot from adult zebrafish brain. Black triangle outlines high frequency contact region between the *meis1b* promoter and the highly conserved non-coding region that contains conserved sequence blocks from the human *LINC01798*. Note the conserved regions overlap ATAC and enhancer (H3K27ac) peaks. B’. zoomed view of shared promoter region between *meis1b* and the antisense transcript si:dkey-196j8.2. Orange shaded bar marks a signature of a shared divergent promoter (ATACseq peak between H3K27ac/H3K4me3 peaks) between *meis1b* and si:dkey-196j8.2.

We examined the promoter for conserved chromatin architecture because our Capture-C data showed the insomnia variant of interest interacts with the promoters of *MEIS1* and *MEIS1-AS3*. Markers of accessible, active promoters (H3K4me3 + H3K27ac + ATAC-seq peaks) were found in the promoter regions of both *MEIS1* (**Figure 3A and A’**) and *meis1b* (**Figure 3B and B’**). Interestingly, an antisense transcript, *si:dkey-196j8.2*, is directly upstream of the promoter of *meis1b* in zebrafish (**Figure B and B’**), similar to the human *MEIS1:MEIS1-AS3.* The presence of double peak H3K27ac/H3K4me3 histone marks, together with ATAC-seq peaks near the transcription start sites of *MEIS1-AS3* and *MEIS1* are indicative of a divergent shared promoter driving transcription in both directions^63,64^ (**Figure A’**). This same chromatin signature was observed between *meis1b* and *si:dkey196j8.2* (**Figure B’**). In contrast, no upstream antisense transcript shares a promoter with *meis1a* (**Figure S4**), and the promoter region of *meis1a* only shows active marks of transcription at the early embryo stage in zebrafish and not in adult zebrafish brain (**Figure S4**).

rs13033745 also showed multiple distal contacts spanning the *MEIS1* TAD; therefore, we examined a comparable region surrounding the zebrafish *meis1* ohnologs for similar chromatin signatures. rs13033745 forms a chromatin loop with the accessible promoter of the long non-coding RNA, *LINC01798*, which is a conserved sequence fragment in zebrafish (**Figure 3A**). *LINC01798* is not annotated in the zebrafish, so we performed a BLAST search against the zebrafish genome and observed that the region maps to the 3’ region distal to *meis1b* (**Figure 3B**) with no comparable sequence homology near *meis1a* (**Figure S4**). The zebrafish transcript downstream of *meis1b*, CR788226.2 (ENSDART00000179158.1), has a continuous 330bp region with 83.9% sequence identity to the human *LINC01798* (**Figure S5**). Notably, these HCNEs show histone marks of active enhancers (H3K27ac + ATACseq peaks) in zebrafish (**Figure 3B**). Similar to human, these distal HCNEs form chromatin contacts with the promoter of *meis1b* as indicated by Hi-C from zebrafish brain^53^ (**Figure 3B**). As this downstream intergenic region has been shown in zebrafish to harbor PAX6 binding sites that modulate *meis1* expression^60^, we overlapped our chromatin Capture C loops with predicted conserved PAX6 binding sites. Indeed, this promoter-interacting loop overlaps a conserved PAX6 binding site (**Figure 3A**), suggesting a gene regulatory interaction between these two hindbrain-expressed developmental transcription factors that may be impacted by rs13033745.

Taking these observations in both the public domain and our generated datasets together, we conclude that the chromatin landscape and non-coding sequence conservation of the region surrounding *meis1b*, the ohnolog which produced the insomnia-like phenotype, closely resembles that of the regulatory region surrounding the human *MEIS1* gene. The conservation of cis-regulatory architecture surrounding *MEIS1* underscores the importance of coordinated gene control at this locus.

### The MEIS1 locus is implicated in cerebellar granule cell development across species

To next determine a potential cellular mechanism through which *MEIS1* could influence sleep, we queried public gene expression databases^54^. Bulk tissue eQTL data from GTEx indicate that the putatively causal insomnia GWAS risk variant we implicated, rs13033745(G), colocalizes with an eQTL that increases *MEIS1-AS3* expression in the cerebellum (**Figure 4A**). *MEIS1* and *MEIS1-AS3* are oriented in a non-overlapping head-to-head position with *MEIS1-AS3* being transcribed from the reverse strand (**see Figure 3A’**). Expression of antisense transcripts in this orientation are suspected to control cell-specific sense transcriptional regulation through shared promoter elements^63,65,66^ (see **Figure 5**). If so, one would hypothesize that *MEIS1-AS3* expression is coordinated with *MEIS1* in a subset of cells^67^. We reanalyzed public scRNAseq data from human cerebellum^54^ indicating that *MEIS1* is highly expressed in the developing cerebellum, predominantly within *NEUROD6/GABRA2+* progenitors that give rise to *PAX6/GABRA6+* cerebellar granule cells (**Figure 4B-C**). Analysis of the non-coding transcripts in contact with the *MEIS1* promoter, *MEIS1-AS3* and *LINC01798*, show *MEIS1-AS3* is most highly expressed in *NEUROD6/GABRA2+* glutamatergic deep nuclei (glutDN) and other cell types along the nuclear transitory zone (NTZ) including granule cell progenitors (**Figure 4B-C and Figure S6**). However, its expression is limited in *PAX6/GABRA6+* mature granule cells (**Figure 4B-C and Figure S6B**). Conversely, *LINC01798* is most highly expressed in maturing *PAX6+* cerebellar granule cells (**Figure 4B-C and Figure S6C**). *LINC01798* is spatio-temporally coordinated with both *MEIS1* and *PAX6* throughout cerebellar development (**Figure 4C and Figure S6**), supporting the finding that PAX6 binding sites are observed within *MEIS1-*promoter-interacting region of *LINC01798*.

**Figure 4.**
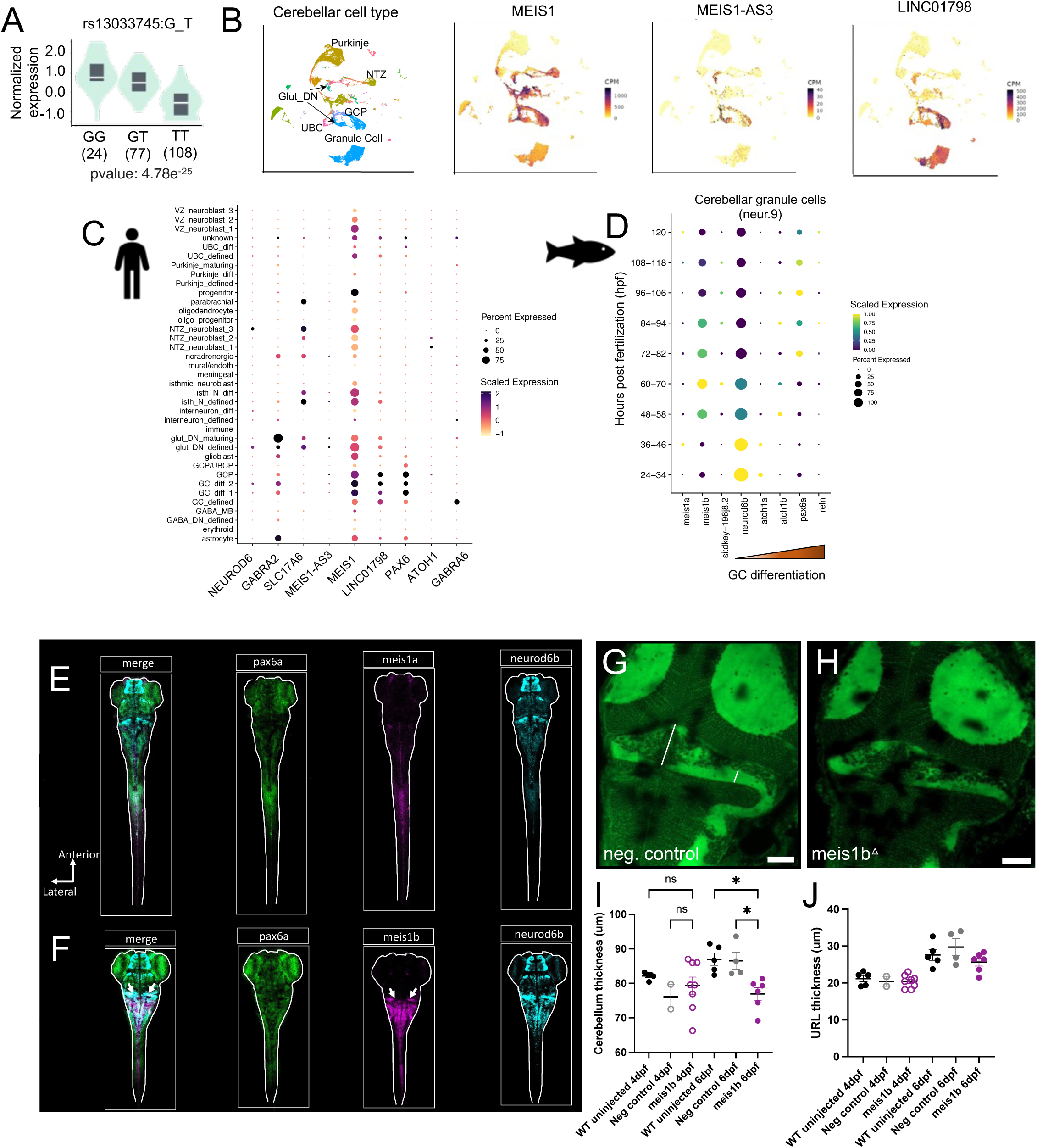
**Conservation of *MEIS1* gene expression across cerebellar granule lineage**. A. Cerebellar eQTL for *MEIS1-AS3* from human GTEx data for the insomnia variant rs13033745. The G allele is associated with increased risk of insomnia and increased expression of *MEIS1-AS3*. B. UMAP plots of scRNAseq data collected from human cerebellum^54^ showing *MEIS1*, *MEIS1-AS3*, and *LINC01798* expression is restricted to the glutamatergic granule cell lineage. C. scRNAseq data from human cerebellum^54^ showing pattern of expression for *MEIS1*, *MEIS1-AS3*, and *LINC01798* with markers of early granule cells (*NEUROD6*, *GABRA2*, and *SLC17A6*) and mature granule cells (*PAX6*, *ATOH1*, and *GABRA6*). D. scRNAseq data from zebrafish cerebellar granule cells across larval development^55^. Known markers of cerebellar granule cells were selected to examine coexpression pattern with *meis1a*, *meis1b*, and *si:dkey196j8.2*. Markers *neurod6b*, *atoh1a*, *atoh1b*, *pax6a*, and *reln* are shown from left to right to represent their temporal expression across granule cell differentiation. Representative images of whole mount in situ HCR in a wild-type 4dpf larval zebrafish showing localization of *meis1a* (E) and *meis1b* (F) along with early granule cell marker *neurod6b* and mature marker *pax6a*. White arrows indicate granule cell population in F. Confocal images were taken using the same parameters for each marker from the dorsal side using a 5x objective. Auto-scaling was used for image enhancement. Images are displayed for a single z-slice (**Supplementary Videos 1-2** show full z-stacks). BODIPY^493/503^ staining in a 6dpf negative control (G) and *meis1b* crispant (H) zebrafish showing clear outline of the cerebellum and upper rhombic lip (URL). White lines in (G) show where measurements were taken for the cerebellar thickness (middle) and URL thickness (right). I. Cerebellum thickness in uninjected wild-type (WT), negative control-injected, and *meis1b* crispant siblings at 4 and 6dpf. J. URL thickness in uninjected wild-type (WT), negative control-injected, and *meis1b* crispant siblings at 4 and 6dpf. Asterisks indicate Sidak multiple comparison corrected *P*-value following one-way ANOVA within each timepoint (4 or 6dpf). Scale bar is 50𝜇m.

**Figure 5.**
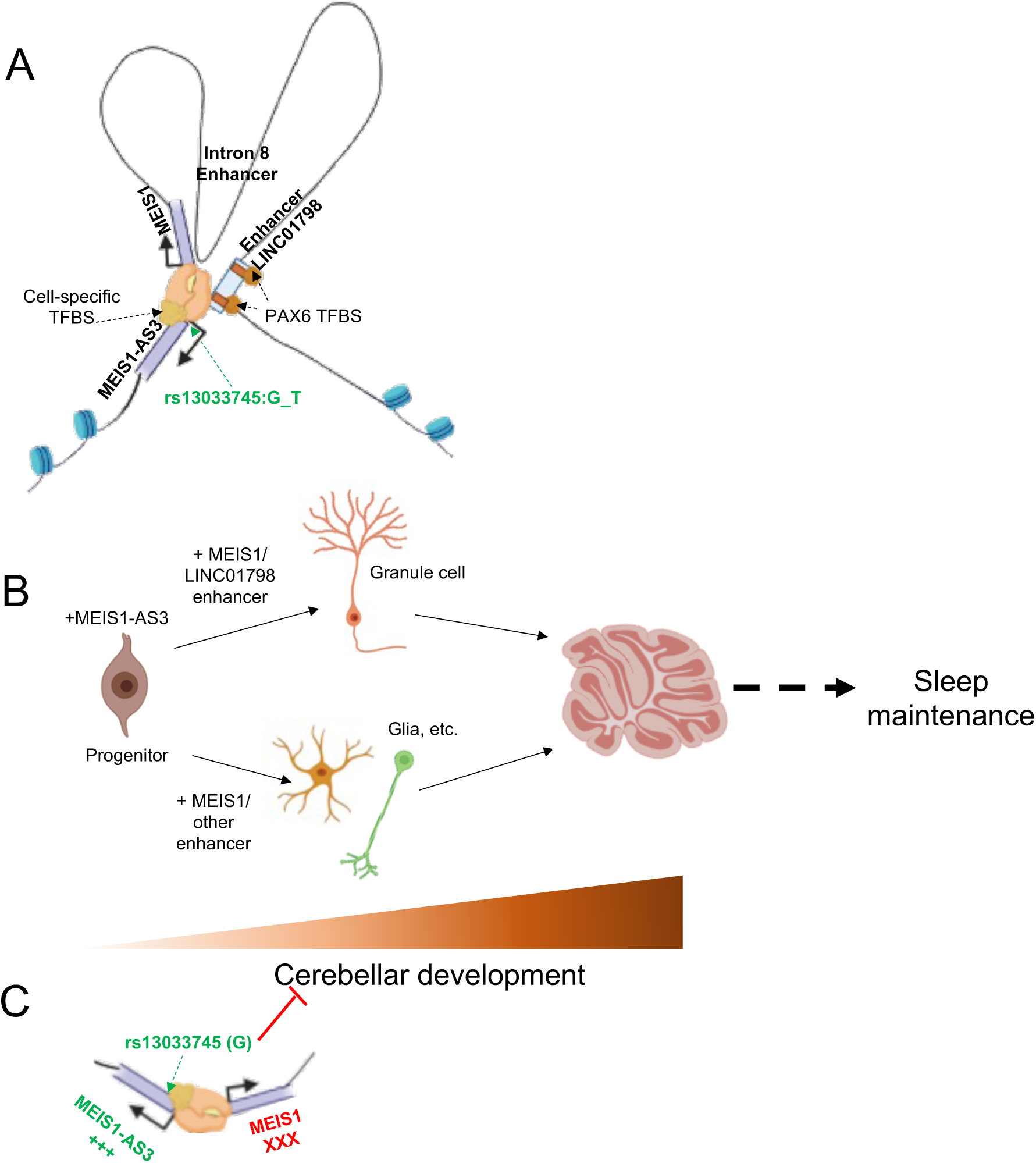
Proposed mechanism of the insomnia associated variant (G) at rs13033745. A. Schematic of the predicted chromatin architecture at the *MEIS1* locus based on data from Figures 2-3. rs13033745 falls in a shared divergent promoter region between *MEIS1* and *MEIS-AS3* Interactions at this promoter region were found between other enhancer-like elements within the introns of *MEIS1* and the *LINC01798* promoter downstream of *MEIS1*. Specifically, distal loops overlap predicted PAX6 transcription factor binding sites. PAX6 is coexpressed with MEIS1 in cerebellar granule cells during maturation. B. Schematic of proposed cerebellar cell differentiation dependent on differential expression of non-coding RNAs based on single cell expression data from Figure 4. *MEIS1-AS3* is highest in progenitor populations of the cerebellum possibly to maintain progenitor pool. Addition of *MEIS1* and cell-specific enhancers would drive differentiation to different cellular populations. *LINC01798* is highest in mature granule cells supporting its role as an enhancer to *MEIS1* for granule cell maturation. C. The insomnia risk allele rs13033745 (G) is associated with increased *MEIS1-AS3* expression in the cerebellum (Figure 4A); therefore, it may increase expression of *MEIS1-AS3* at the expense of *MEIS1* by shifting the divergent promoter to favor *MEIS1-AS3*. This shift toward *MEIS1-AS3* should prevent cerebellar granule cell maturation leading to impaired cerebellar development and deficits in sleep maintenance.

To determine whether *MEIS1* and *meis1b* are expressed in similar populations of cerebellar cells, we subsequently queried public domain scRNAseq data for known markers of the glutamatergic granule cell lineage in zebrafish^68,69^. Similar to humans, we observed *meis1b* expression peaks in *neurod6b+/pax6a+* granule cells early in development (60-70hpf) aligning with cerebellar granule cell differentiation^68,69^ (**Figure 4D and Figure S7**). In addition, similar to human *MEIS1-AS3*, the zebrafish antisense transcript, *si:dkey-196j8.2,* is expressed in a subset of early progenitors that give rise to granule cells (**Figure S7**). While *si:dkey-196j8.2* shows coordinated expression with *meis1b* in some cell types, it shows distinct expression in two hindbrain/rhombic lip progenitor populations (neur6 and neur39) (**Figure S7**). *Meis1a* expression was very low in these populations (**Figure 4D and Figure S7**).

To confirm colocalization of *meis1b* with these common markers of granule cell development *in vivo* and identify cell populations that could be vulnerable to *meis1a/b* loss, we performed *in situ* hybridization chain reaction (HCR)^47,48^ on whole-mount larval zebrafish at 4dpf, representing a timepoint when the cerebellar circuitry is intact but differentiation is occurring^68^. We observed that *meis1a* is only expressed at a very low level at 4dpf in the zebrafish (**Figure 4E, Supplementary Video 1**) and is predominantly localized to the spinal cord, consistent with the single cell expression data (**Figure 4D**). *Meis1b* colocalized with *neurod6b* in the cerebellum at 4dpf and to a lesser degree with *pax6a* (**Figure 4F, Supplementary Video 2**), suggesting *meis1b* is in a granule cell-like progenitor population at this timepoint.

Given we observed the sleep abnormalities beginning at day 6 post fertilization in *meis1b* loss-of-function crispants, and cerebellar differentiation occurs between 4 and 6dpf^68^, we measured the size of the cerebellum at 4 and 6dpf using a Bodipy dye which labels the cerebellar neuropil providing a clear outline of the cerebellum and rhombic lip^68,70^ (**Figure 4G-H**). At 4dpf, we found no significant difference in cerebellar or upper rhombic lip width (**Figure 4I-J**). However, at 6dpf, the cerebellum was significantly smaller in *meis1b* crispants compared to negative controls or sibling-matched wild-type fish that did not receive an injection (**Figure 4I**). The upper rhombic lip, which is where granule cells originate, did not differ in width (**Figure 4J**).

Together, these data provide support for MEIS1 in the maturation of the cerebellum across species and indicate the non-coding genomic architecture at the *MEIS1* locus may underlie the fragmented sleep phenotype through its influence on *MEIS1* expression during cerebellar development.

## Discussion

Despite abundant GWAS data for complex traits, functional validation within a living model system is often not performed. Here, we demonstrate an approach for moving from GWAS-implicated effector genes to validation in model organisms and integrate publicly accessible data with our human cell-based variant-to-gene approach to propose and test a genetic mechanism contributing to insomnia.

There is a need for preclinical models of genetic disorders to allow for therapeutic development, but a scalable *in vivo* model with human disease-relevance is lacking. Advances in high-efficiency gene editing within zebrafish^40,71^ and scalable sleep phenotyping^42^ allowed us to test multiple candidate genes. In our zebrafish screen, *skiv2l* crispants displayed markedly reduced sleep during the day pointing to a role for this immunomodulatory gene in sleep regulation. Meanwhile, loss of the transcription factor involved in cell fate specification, tcf12, reduced sleep episodes at night. Of our top six candidate genes, CRISPR mutation of *meis1b* produced a phenotype with impaired sleep maintenance and delayed sleep latency, supporting years of GWAS implicating *MEIS1* as a top candidate insomnia risk gene^4,6,7^. *MEIS1* encodes a highly conserved transcription factor involved in developmental patterning and cerebellar development^33,58,59,72,73^, yet its role in sleep regulation is not well understood. In mouse models, *Meis1* deficiency was associated with circadian hyperactivity^32,74,75^, and an RLS-associated point mutation altered sleep consolidation^76^, but phenotypes have been inconsistent. Moreover, it is unclear how these findings translate to sleep maintenance given mice do not consolidate sleep to a single phase. Because zebrafish, like humans, are diurnal with sleep consolidated to the night, we were able to identify a phenotype similar to insomnia in humans (e.g. marked by fragmented sleep restricted to the night) that implies MEIS1 is acting in a diurnal pattern to alter sleep consolidation. These mutants also had an increase in nighttime sleep latency, a common characteristic of insomnia. Our video-tracking assay provides highly sensitive movement data using pixel displacement, which allows us to detect small body movements that may represent an RLS-like phenotype. Using this method, we showed that loss of *meis1b* in fish does not alter nighttime waking activity, arousability, or depth of sleep. Rather *meis1b* mutants transition between wake and sleep more frequently. This represents an important distinction given the cerebellum, which we showed is developmentally impaired in *meis1b* crispants, has been implicated in sleep-wake transitions^77,78^; results point to a key role for this region in sleep maintenance rather than depth. The cerebellum has been largely ignored for its role in insomnia even though insomnia and sleep health GWAS have repeatedly shown the cerebellum is the most enriched tissue for risk gene expression^6,7,79,80^. Thus, we believe our *meis1b* zebrafish model represents an important biological model for sleep fragmentation that is relevant to human disorders.

The broader *MEIS1* region has been shown in multiple species to harbor enhancers that control *MEIS1* expression in the brain^32,60,81,82^. Enhancers in the region encompassing *meis1b* on zebrafish chromosome 13 have also been shown to be expressed in the developing hindbrain and regulate *MEIS1* expression in the retina through PAX6 recruitment^60^. The chromatin interaction we observed with rs13033745 in the *MEIS1-AS3* promoter has not been previously described. A gene-spanning contact has been reported previously for multiple different cell types^37^, indicating distal control of the *MEIS1* promoter region is imperative to its function. Our promoter-focused capture C data also show long-range chromatin contact loops between the *MEIS1/MEIS1-AS3* promoter and the *LINC01798* region that harbors PAX6 binding sites. Given the highly concordant expression pattern of *LINC01798* and *PAX6* in cerebellar granule cells, and the coordination between MEIS1 and PAX6 in cerebellar development^59^, this distal interaction may be a key player in cerebellar patterning. The RLS signal in intron 8, tagged by rs113851554, overlaps an annotated brain enhancer as well as eQTLs in the caudate^32,37^; therefore, these SNPs may alter brain development via different cell-specific trajectories (**Figure 5**).

Because rs13033745 is in an accessible promoter-contact loop across many cell types and colocalizes with an eQTL impacting *MEIS1-AS3* expression in the cerebellum, we implicate this SNP as a putatively causal SNP at the *MEIS1* locus for insomnia. We further propose a model (**Figure 5**) whereby the risk allele rs13033745(G) shifts the expression in favor of *MEIS1-AS3* at the expense of *MEIS1* via transcriptional interference since they appear to share a divergent promoter^65,66^. Therefore, we suggest the increase in *MEIS1-AS3* and/or interaction of the *MEIS1* promoter with other distal enhancers is a mechanism that influences *MEIS1* spatio-temporal regulation leading to altered cellular differentiation within the cerebellum. We noted another proxy variant to the 5’ signal resides in a CTCF site for the sub-TAD. While our approach combining ATACseq and chromatin capture to nominate variants in direct contact with gene promoters did not implicate that site, this may warrant further investigation given the role of CTCF in stabilizing sub-TADs^83^.

We showed that the non-coding architecture is highly conserved only for one ortholog, *meis1b*, in zebrafish, and that this high degree of non-coding conservation correlates with the observed behavioral phenotype. The non-coding conservation around *meis1b* may help determine the cellular and functional specificity of this ohnolog. *MEIS1* and *MEIS1-AS3* show coordinated, but distinct gene expression indicating *MEIS-AS3* transcription is not likely due to transcriptional noise. Recent work has demonstrated divergent antisense transcription is a common feature of lineage-determining transcription factors^63^. Thus, this conserved antisense element is likely critical for regulating *MEIS1* expression during lineage progression of granule cells and/or other progenitors. Indeed, we found that loss of *meis1b* impairs developmental progression of the cerebellum, while others have found loss of *meis1b* impairs retinal specification and tectum size in the zebrafish^73^. Further work will be needed to characterize how these non-coding elements restrict *MEIS1* expression across development.

## Limitations

Previous work has shown CRISPR-mediated deletion of non-coding enhancer-like sequences at the *MEIS1* locus reduces *MEIS1* expression^37^ and that the intron 8 risk allele is associated with reduced *MEIS1* levels^31^; therefore, CRISPR loss-of-function mutations are relevant to the phenotype. However, since the 5’ variant of interest resides in a divergent promoter between *MEIS1* and the corresponding anti-sense RNA across species, we could not reliably manipulate the anti-sense promoter without also disrupting the sense promoter. *MEIS1-AS3* is not highly conserved at the sequence level, rather the orientation of the transcript with respect to the sense RNA is conserved as has been shown for constrained cis-regulatory elements^29^. Anti-sense RNAs are often not conserved at the sequence level across species, instead they act through transcriptional interference^66^. While possible, it is unlikely the upstream anti-sense RNA is acting in *trans* to suppress *MEIS1* expression as it has very low sequence complementarity to *MEIS1* and does not overlap *MEIS1* transcripts, thus deletion of the exonic regions of the anti-sense transcript may yield little-to-no effect on *MEIS1* itself. Future work performing targeted genetic manipulation will help elucidate the role of the anti-sense RNA.

While *MEIS1* is primarily restricted to excitatory granule cell lineage cells, we cannot rule out the downstream effects that may influence other cell types such as Purkinje cells. Granule cells are important for the development of Purkinje cells^84^; therefore, the sleep phenotype could be due to disruption of either cell population. *Meis1b* is most highly expressed in the hindbrain and cerebellum in zebrafish, and its loss impaired cerebellar development; however, *meis1b* expression in other cell types may also contribute to the sleep phenotype observed. Additional experiments will be needed to determine the precise cell type responsible for the sleep fragmentation.

By selectively targeting *meis1b* with CRISPR-Cas9, we could examine the contribution of the ohnolog most highly expressed in the developing hindbrain, without impacting *meis1a*, which is expressed in the developing somite. This allowed us to mitigate potential developmental defects due to broad loss of *meis1* expression and provide greater cellular specificity to our genetic manipulation. Loss of *meis1a* alone did not produce a significant sleep phenotype, and *meis1a* is not highly expressed in the brain; therefore, compensation from *meis1a* is not expected to play a role in the single *meis1b* mutants. Previous work has also confirmed *meis1a* does not appear to compensate for *meis1b*^85^. It is, however, possible that duplicated ohnologs may compensate to some degree, which may explain why several of the other genes we targeted (*cbx1a/b* and *gnb3a/b* had modest effects on sleep).

## Conclusion

The comprehensive intersection of human spatial genomics paired with *in vivo* phenotyping and genomic conservation analyses has allowed us to reveal a novel association for *MEIS1* with insomnia. We show that functional phenotyping in a simple vertebrate is critical and insightful, pointing to deeply conserved non-coding control of regulatory genes. This type of conserved non-coding regulation is not unique to the *MEIS1* locus and may explain many of the non-coding GWAS signals associated with complex traits.

### Data and code availability

Full datasets containing sleep/activity data are available at http://datadryad.org/share/I07vjxcKg3i5Atl_H1A60C02e3A-gFLA2bLXofNpC7E. MATLAB scripts and full protocol for sleep/activity analysis can be found at DOI: <u>10.21769/BioProtoc.4313.</u>

## Acknowledgements

The work was supported by NIH grant T32 HL07953, T32 HL170968, R01 HL143790, P01 HL094307, and P01 HL160471. Dr. Chesi is supported by R35 HG011959 and R01 NS135075. Dr. Dr. Veatch is supported by P20 GM130423. Grant is supported by NIH awards R01 AG057516 and R01 HD056465 and the Daniel B. Burke Endowed Chair. Authors would like to thank the CHOP Zebrafish Core for assistance in zebrafish care, maintenance, and CRISPR experimentation. Authors thank the UPenn CDB microscopy core for assistance with image acquisition. Authors are also grateful to Dr. Jason Rihel and Joshua Donnelly for their training and assistance in HCR protocol and probe design.

## Authorship Contribution

A.J. Z. Designed and conducted experiments, analyzed data, and wrote the manuscript. E.A., M.C.P., F.D.B., B.T.K., and P.Z.L. conducted experiments and analyzed data. Z.Y.S and T.R.T. conducted experiments. J.P. and A.K. assisted in gene prioritization and conceptualization. J.A.P, A.D.W, and O.J.V conceptualized experiments. A.C. conceptualized experiments and acquired and analyzed capture C datasets. P.R.G., A.C.K., A.I.P, and S.F.A.G conceptualized experiments, coordinated and managed collaborations, and supervised the study. All authors reviewed and edited the manuscript.

## Web Resources

**DIOPT** https://www.flyrnai.org/cgi-bin/DRSC_orthologs.pl

**ensembl** https://useast.ensembl.org/

**UCSC** genome browser https://genome.ucsc.edu/

**WashU epigenome** browser https://epigenomegateway.wustl.edu/

**3D genome browser** https://3dgenome.fsm.northwestern.edu/

**Kaessman Human Cerebellar Gene Expression Portal** https://apps.kaessmannlab.org/sc-cerebellum-transcriptome/

**DanioCell** https://daniocell.nichd.nih.gov/

**LD matrix** https://ldlink.nih.gov/?tab=ldmatrix

