## SupplementaryFigures for "Cross-species variant-to-function analyses implicate *MEIS1* in conferring sleep abnormalities and impaired cerebellar development"

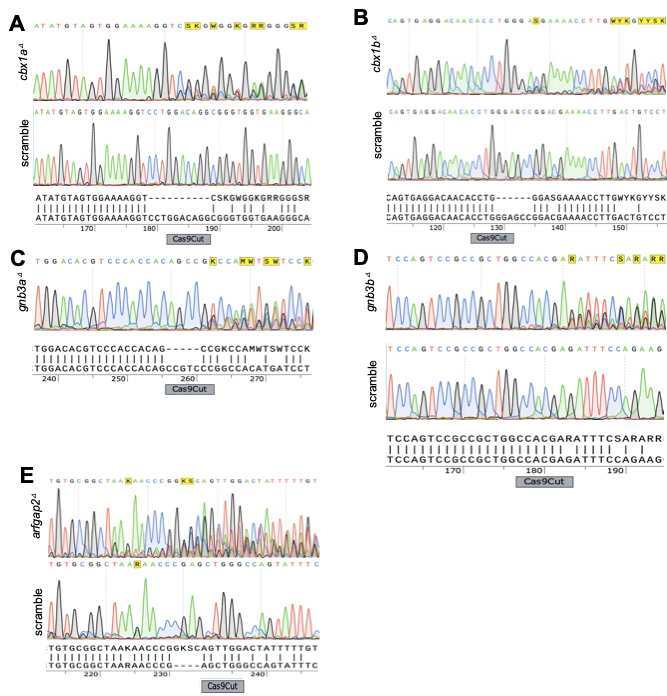
Figure S1

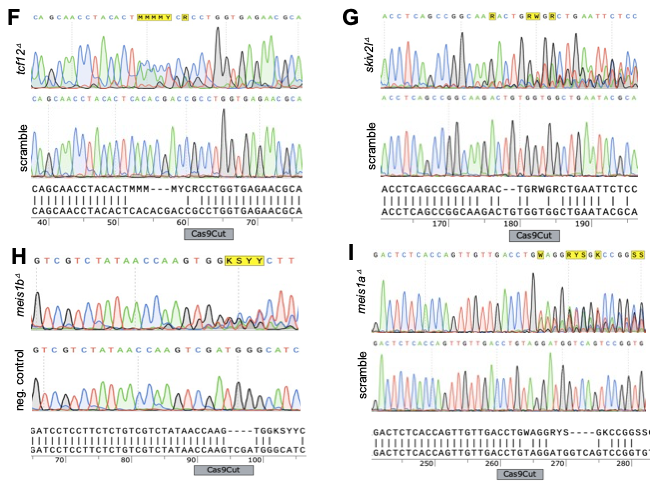

Figure S2

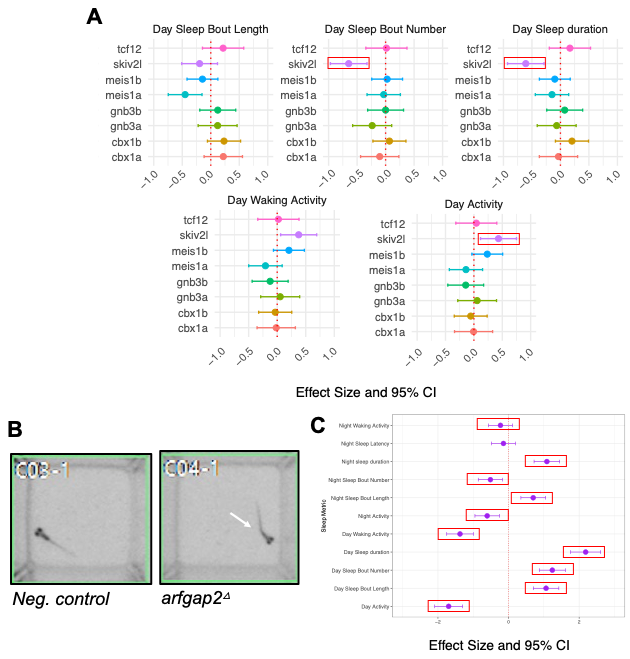

Figure S3

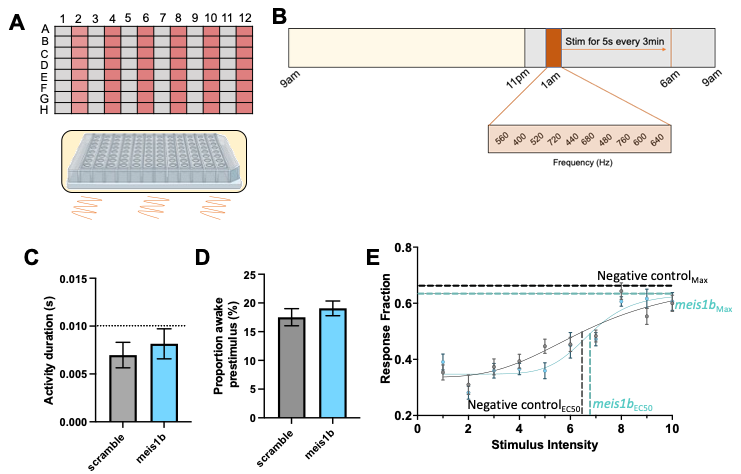

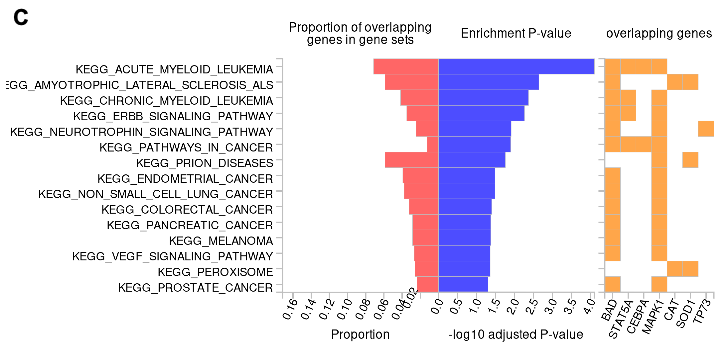

Figure S4

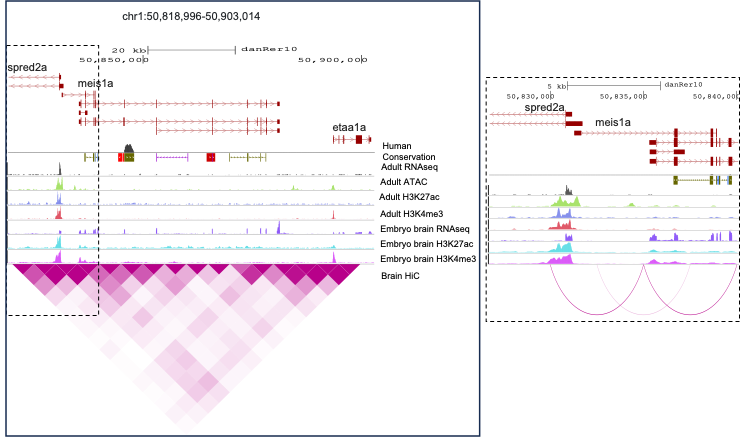

Figure S5

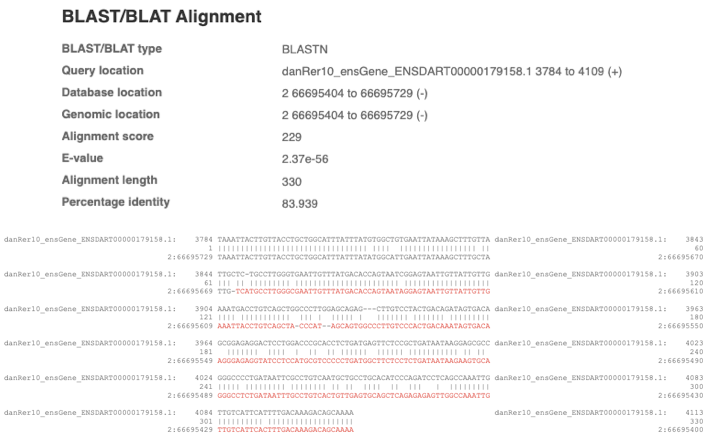

Figure S6

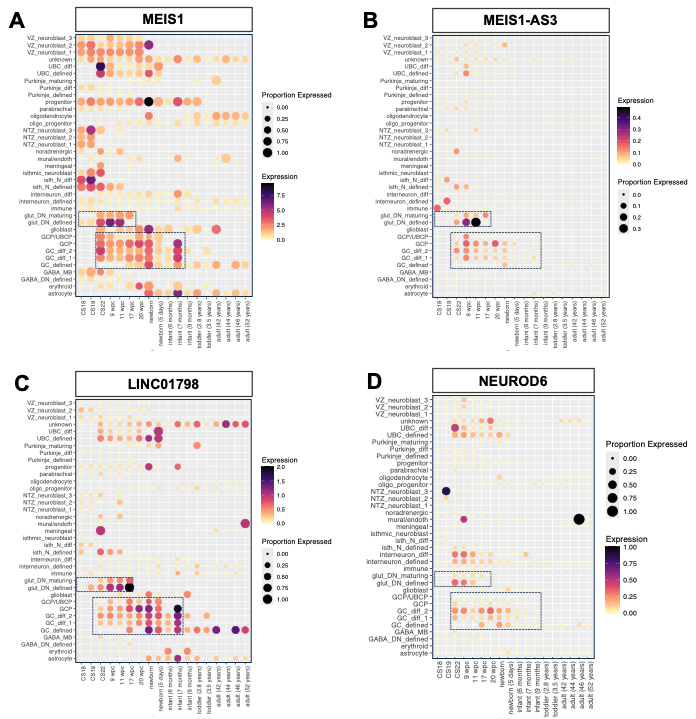

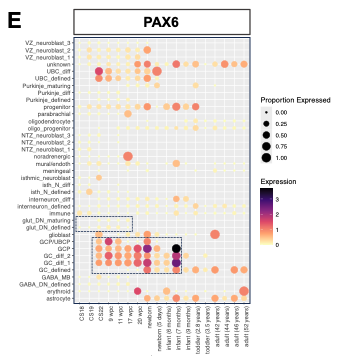

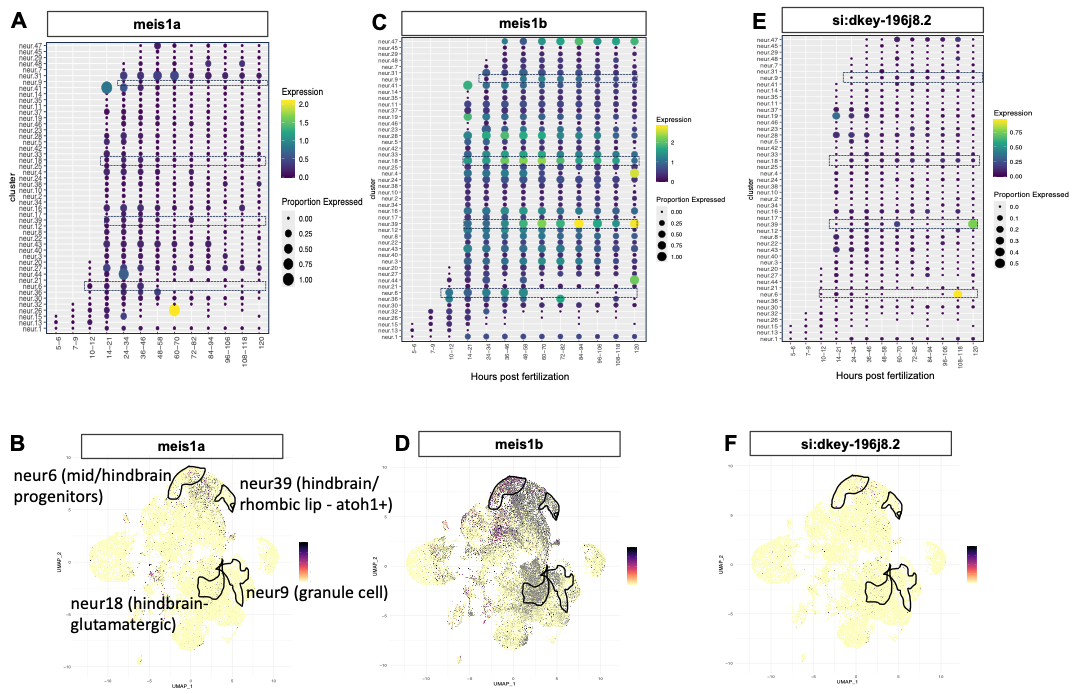
Figure S7

**Figure S1. Confirmation of CRISPR/Cas9 on-target mutation.** A-I. Representative Sanger electropherograms aligned with sequence and Cas9 cut site for each candidate gene. Traces are shown for a single crispant and negative control (scramble-injected) sibling from each gene group. Sequence alignment shows consequence of the cut that shifts the reading frame in all cases due to non-multiple of 3bp changes.

**Figure S2. Additional sleep-wake metrics in crispant fish.** A. Data for *tcf12*, *skiv2l*, *meis1a*, *meis1b*, *gnb3a*, *gnb3b*, *cbx1a*, and *cbx1b* crispants. Data are presented as effect size estimate (standardized mean difference) +/- 95% confidence interval (CI) for each crispant group versus its sibling scramble-injected controls. B. Example of *arfgap2* crispant from sleep assay showing tail curvature. C. Data for *arfgap2* crispants presented as effect size estimate $\pm$ 95% confidence interval (CI) for *argap2* crispants versus sibling scramble-injected control. N for each experiment: *skiv2l* (80 crispants/ 77 controls), *meis1a* (94 crispants/ 86 controls), *meis1b* (111 crispants/ 100 controls), *gnb3a* (68 crispants/64 controls), *gnb3b* (75 crispant/ 80 controls), *cbx1a* (60 crispant/ 79 controls), *cbx1b* (102 crispant/ 83 controls), (74 crispant, 49 controls), *arfgap2* (58 crispant/79 controls). Red boxes indicate measures that reached significance following correction as described in **Methods**.

**Figure S3. Arousal threshold is not altered in *meis1b* crispants.** A Schematic of plate design for arousal threshold assay where crispants and controls are placed into alternating columns of a 96-well plate which is placed atop a vibration unit within the standard sound-and light-attenuating Zebrabox (ViewPoint Life Sciences) B. Ten different mechanoacoustic stimulus intensities are pseudorandomly presented for 5 seconds at a time every three minutes from 1am to 6am during the lights off period (ZT16-21). C. Total activity duration during the 5-second prestimulus window per fish (t_(59)_ = 1.16, *P* = 0.25). All fish fall below the threshold (dotted line) for sleep (0.01s/5sec). D. Proportion of fish awake in the 5-second prestimulus window (activity>0.01s/5sec) across all stimuli presentations (t_(198)_ = 1.56, *P* = 0.12). E. Stimulus response curve plotting the proportion of sleeping fish that were aroused by each stimulus intensity. No significant difference observed for EC_50_ between groups: F_(1, 182)_ = 0.031, *P* = 0.86. Maximum response and EC_50_ are shown by dashed lines. EC₅₀ represents the stimulus intensity required to elicit an arousal response that is 50% of the maximum response rate observed in the group. N = 29 controls and 32 *meis1b* crispants.

**Figure S4. WashU epigenome browser plot showing chromatin landscape at the *meis1a* locus.** Chromosomal coordinates are shown for the danRer10 genome build. Conservation is shown with reference to the human genome with human chain-net track colored by chromosome. Chromatin tracks show data from Yang et al. for RNAseq, ATACseq, H3K27ac, and H3K4me3 marks from zebrafish adult brain and embryo along with a HiC interaction plot from adult brain. Inset shows promoter region (black dashed box) of *meis1a*.

**Figure S5.** BLAST alignment of the zebrafish CR788226.2 genomic sequence to the human genome.

**Figure S6.** **Gene expression of *MEIS1*-associated genes in the developing human cerebellum.** Single cell RNAseq data from Sepp et al., 2024 were analyzed to show *MEIS1* (A), and the non-coding RNAs that form chromatin contacts with the risk variant rs13033745, *MEIS1-AS3* (B), *LINC01798* (C), as well as a marker of glutamatergic deep nuclei/granule cell progenitors (*NEUROD6*) (D) and maturing granule cells (*PAX6*) (E). Color represents scaled expression value and dot size represents proportion of cells expressing the transcript >0. Data are presented for cell type and stage of development from Carnegie Stage 18 to adulthood. Black dashed boxes outline glutamatergic/granule-lineage cell types and time points with coordinated expression.

**Figure S7. Gene expression of *meis1a/b* and antisense RNA *si:dkey196j8.2*.** Single cell RNAseq data analyzed from DanioCell**.** Expression of *meis1a* (A-B), meis1b (C-D), and *si:dkey196j8.2* (E-F) across different neuronal cell types and developmental time points from 0 -120 hours post fertilization. Color represents scaled expression value and dot size represents proportion of cells expressing the transcript >0. Clusters of interest in A,C and E, are outlined in black in B,D, and F UMAP plots.

**Supplementary Video 1**

Confocal z-stack through dorsally-mounted 4dpf zebrafish brain following HCR. Magenta = *meis1a*, cyan = *neurod6b*, green = *pax6a*. Autoscaling was applied to the stack to enhance brightness. Image was acquired using 10x objective with a 1$\mu$m z-step.

**Supplementary Video 2**

Confocal z-stack through dorsally-mounted 4dpf zebrafish brain following HCR. Magenta = *meis1b*, cyan = *neurod6b*, green = *pax6a*. Autoscaling was applied to the stack to enhance brightness. Image was acquired using 10x objective with a 1$\mu$m z-step.
