## Supplementary material for "Cross-species variant-to-function analyses implicate *MEIS1* in conferring sleep abnormalities and impaired cerebellar development": Tables1-2

| **Phenotype** | **Mean (95% CI)** | | **Difference**  **(95% CI)** | ***P*-value** |
| --- | --- | --- | --- | --- |
|  | **Neg. control** | ***meis1a^△^*** |  |  |
| **Night** | | | | |
| Hourly Sleep, minutes | 37.42 (35.28, 39.56) | 38.04 (36.18, 39.91) | 0.63 (-2.18, 3.43) | 0.9043 |
| Hourly Sleep Bouts, number | 7.45 (7.02, 7.88) | 7.8 (7.37, 8.23) | 0.35 (-0.25, 0.96) | 0.1312 |
| Sleep Latency, minutes | 4.99 (4.37, 5.62) | 4.34 (3.71, 4.96) | -0.66 (-1.53, 0.22) | 0.0303 |
| Bout Length, minutes | 6.08 (5.05, 7.12) | 6.59 (5.30, 7.88) | 0.51 (-1.14, 2.15) | 0.7724 |
| Hourly Activity, sec | 28.96 (26.22, 31.69) | 27.66 (25.49, 29.83) | -1.30 (-4.74, 2.14) | 0.8658 |
| Avg. Wake Act., sec/awake min | 0.69 (0.65, 0.74) | 0.66 (0.63, 0.70) | -0.03 (-0.08, 0.03) | 0.7225 |
| **Day** | | | | |
| Hourly Sleep, minutes | 10.42 (8.74, 12.10) | 9.32 (7.91, 10.74) | -1.09 (-3.26, 1.08) | 0.3100 |
| Hourly Sleep Bouts, number | 3.83 (3.36, 4.30) | 3.76 (3.31, 4.21) | -0.07 (-0.72, 0.58) | 0.6644 |
| Bout Length, minutes | 2.79 (2.51, 3.06) | 2.34 (2.22, 2.46) | -0.45 (-0.75, -0.15) | 0.0129 |
| Hourly Activity, sec | 144.28 (134.15, 154.40) | 138.03 (129.45, 146.62) | -6.24 (-19.35, 6.87) | 0.3476 |
| Avg. Wake Act., sec/awake min | 2.57 (2.41, 2.74) | 2.42 (2.28, 2.56) | -0.15 (-0.36, 0.06) | 0.1728 |
| Statistically significant differences after Hochberg step-up correction are indicated with an asterisk. Mean and 95% confidence interval. Mean difference and 95% CI | | | | |

**Table 1. Sleep parameters in *meis1a* crispant and negative control fish.**

| **Phenotype** | **Mean (95% CI)** | | **Difference**  **(95% CI)** | ***P*-value** |
| --- | --- | --- | --- | --- |
|  | **Neg. control** | ***meis1b^△^*** |  |  |
| **Night** | | | | |
| Hourly Sleep, minutes | 38.32 (36.38, 40.26) | 36.93 (35.25, 38.61) | -1.39 (-3.93, 1.15) | 0.3015 |
| Hourly Sleep Bouts, number | 7.19 (6.79, 7.59) | 8.53 (8.16, 8.89) | 1.34 (0.80, 1.87) | <0.0001* |
| Sleep Latency, minutes | 4.52 (4.07, 4.97) | 5.65 (5.13, 6.17) | 1.13 (0.44, 1.82) | 0.0004* |
| Bout Length, minutes | 6.55 (5.70, 7.41) | 5.06 (4.48, 5.65) | -1.49 (-2.52, -0.46) | 0.0024* |
| Hourly Activity, sec | 29.59 (26.74, 32.44) | 30.95 (28.70, 33.21) | 1.37 (-2.21, 4.94) | 0.2532 |
| Avg. Wake Act., sec/awake min | 0.69 (0.65, 0.74) | 0.69 (0.65, 0.73) | 0.00 (-0.06, 0.06) | 0.7212 |
| **Day** | | | | |
| Hourly Sleep, minutes | 9.77 (8.37, 11.17) | 9.06 (7.69, 10.44) | -0.71 (-2.66, 1.24) | 0.1625 |
| Hourly Sleep Bouts, number | 4.02 (3.53, 4.50) | 4.10 (3.44, 4.76) | 0.08 (-0.73, 0.90) | 0.2695 |
| Bout Length, minutes | 2.49 (2.33, 2.65) | 2.37 (2.23, 2.52) | -0.12 (-0.33, 0.10) | 0.2147 |
| Hourly Activity, sec | 121.60 (113.41, 129.78) | 132.52 (122.92, 142.13) | 10.93 (-1.62, 23.48) | 0.1980 |
| Avg. Wake Act., sec/awake min | 2.16 (2.02, 2.29) | 2.32 (2.16, 2.47) | 0.16 (-0.05, 0.37) | 0.2695 |
| Statistically significant differences after Hochberg step-up correction are indicated with an asterisk. Mean and 95% confidence interval. Mean difference and 95% CI | | | | |

**Table 2. Sleep parameters in *meis1b* crispant and negative control fish.**
